# Combining Focal-ERG, Fluorescein Angiography, and SD-OCT for Retinal Neurovascular Analysis in a Mouse OIR-Model

**DOI:** 10.1101/2019.12.12.874271

**Authors:** K.P. Mitton, M. Deshpande, S.C. Wong, E. Guzman, M. Cheng, W. Dailey, R. Schunemann, M. Trese, K. Drenser

## Abstract

The development of non-invasive live ocular imaging and electrophysiological test systems for rodent eyes provides new tools for not only averaged analysis of the entire retina but also the ability to see, test, and compare different subregions of the same retina. These new capabilities provide the possibility for more detailed examinations of local structural and functional relationships within a single eye and the ability to also follow changes longitudinally over time. We have developed protocols based around the Micron-III/IV retinal imaging camera system for combining fluorescent imaging of the neural retinal micro-vasculature by FA (fluorescein angiography), imaging of all neural retinal layers by SD-OCT (Spectral-Domain Ocular Coherence Tomography), and focal “spot” light-targeted electroretinography (Focal-ERG) to relate the local neurovascular unit structure to the inner (photoreceptor) and outer-retinal electrical response to light stimulation. For demonstration purposes we have used the popular mouse oxygen induced retinopathy (OIR) model, which causes radial central patches of retinal neuron loss mostly in zones away from and between the primary retinal arteries and veins. In this model, the loss of central microvasculature is induced developmentally in mouse litters exposed to 75% oxygen from age P7 to P11. Return to room air on P12, causes several days of retinal ischemia during which neurons, mostly of the inner retina, perish. Bipolar and ganglion cell death ends as neovascular growth revascularizes the central retina. This model provides for non-uniform retinal damage as well as gradual progression and resolution over time. The OIR model was used to generate regions of inner retinal neuron loss in B6.Cg-Tg^*Thy1-YFP*^ mice. Using image-guided focal-ERG, the dark-adapted mixed rod-cone light response was compared using stimulation of small circular (0.27 mm diameter) target areas located in the central retinas of the same eyes (OIR and control). The same areas of the same retinas were followed over three ages after revascularization (P21, P28 and P42).

**Conclusions:** Combined FA and SD-OCT imaging can provide local geographic specific information on retinal structural changes and be used to select different retinal areas within the same eye for testing of local light response. This analysis strategy can be employed for studies with rodent disease models that do not uniformly impact the entire retinal area. Combining these techniques would also be useful for testing gene and cell replacement therapies in retinal degeneration models where typically a small zone of the retina is treated. Both treated and untreated retinal zones within the eye can be followed non-invasively over many weeks.

**SUMMARY:** Mouse models utilized for retinal disease research including retinal vascular models can display nonuniform changes over the entire retina. Damage or loss of retinal layers and retinal neurons due to hypoxia can impact some retinal areas while leaving adjacent regions unaltered. Combining vascular imaging by fluoresceine angiography, vascular imaging and retinal layer imaging by SD-OCT, and focal-ERG provides us with new tools to examine retinal structure-function relationships within a single retina.

## INTRODUCTION

While the Human retina is routinely monitored with non-invasive methods, there is a growing awareness that combining multiple imaging modes and targeted functional testing can provide a more complete understanding of both development and regional pathophysiology. In various Human retinal diseases, the initial pathology and the progress of retinal changes are rarely uniform throughout the entire retina area. There are often profound differences in both the radial direction and distance from the disc, or between the central and peripheral retina. Useful disease research models, such as the oxygen induced retinopathy (OIR) model, or testing of sub-retinal injections (genes/cells) would also benefit from the ability to compare different retinal areas within the same eye [1–4]. In this report, we describe the application of non-invasive retinal imaging, combined with focal-ERG, to provide geographical analysis that can also be followed over time after ischemia in the mouse OIR model.

In humans, damage to the post-photoreceptor sensitivity is common in retinopathy of prematurity (ROP) patients depending on the severity of disease (Fulton et al, 2009) [5]. Both photoreceptor sensitivity and post-photoreceptor response are diminished more in severe ROP compared to mild ROP patients, with cone responses less affected than rod responses (Fulton 2008) [6]. Rod photoreceptor function and post-photoreceptor function is also diminished in rat OIR models [7,8]. More recent studies in humans indicate that some ROP patients likely experience recovery of post-photoreceptor function. This is based on the finding that older patients who were classified as having mild ROP as infants have improved post-photoreceptor function compared to infant mild ROP patients [9]. Nakamura et al. (2012) reported an average reduction in the full-field ERG B-wave amplitude from the retinas of OIR mice, consistent with a significant loss of bipolar cells [10]. They also noted a partial recovery of the B-wave amplitude over several weeks post-damage.

Neural retinal maturation and retinal vascular development occurs post-natal in mice. For the oxygen induced retinopathy model, mice are exposed to 75% oxygen for five days, from age P7 to P12. This period starts about the time that the superficial vascular bed is just reaching the peripheral retina, having originated from the optic disc. In the high oxygen environment the normal development of all three vascular beds is impaired and the superficial capillary bed degenerates through apoptosis. After five days of 75% oxygen, by age P12, the central retina is devoid of any capillary beds and thus avascular. Down-regulation of retinal Vegf (Vascular Endothelial Growth Factor) gene expression is a major factor responsible for this vaso-obliteration mechanism [11]. Upon return to room air (21% oxygen), this central retinal zone becomes hypoxic and that results in an increased concentration of VEGFA (vascular endothelial growth factor A) and an aggressive revascularization of the central retina [12]. This process brings oxygen back to the central retina by age P21. Until this recovery to normoxia by age P21, bipolar cell loss can be extensive in the central retina and the thinning of the INL (inner nuclear layer) may be substantial [13].

Since full-field ERG stimulates both the more central OIR damaged retina and the non-damaged peripheral retina, an average retinal ERG is obtained that does not differentiate the two zones. Using focal ERG, a light-spot can be aimed at a descrete small zone of choice and used to compare regions within the same retina. In the mouse OIR model, this provides the potential to test more affected and less severely affected retinal zones, and to follow some recovery of the B-wave response over time. Using fluorescein angiography and SD-OCT, it is possible to find regions of the OIR retinas that have suffered permanent loss of inner retinal neurons and thus select zones for focal-ERG testing.

## PROTOCOL

### Animals

This study was approved by the Oakland University IACUC and complied with the ARVO Statement for the Use of Animals in Ophthalmic and Vision Research. Mice expressing YFP (yellow fluorescent protein) in a subset of ganglion cells (B6.Cg-Tg(Thy1-YFP)HJrs/J) were obtained from the Jackson Laboratory (Bar Harbor, ME).

### Oxygen-induced Retinopathy

Pre-weanling litters were housed in 75% oxygen for five days, as per Smith et al. (1993), from post-natal age P7 to P12. Litters were then returned to room air. This exposure window was late enough to minimize dilation effects on the regressing hyaloid vessels, and early enough to overlap with retinal vascular development. A maximal neovascularization response occurs in this model between age P17 and P21.

### Micron-III system imaging and Focal-ERG

Focal-ERG analysis was completed in the Pediatric Retinal Research Laboratory’s retinal imaging and ERG suite, at the Eye Research Institute of Oakland University. The suite was equipped with variable dim red lighting (4-15 Lux) for working on dark-adapted rodents. Focal-ERG recordings were from dark-adapted mice using a Micron-III camera-mounted focal-ERG system (Phoenix Research Labs, Pleasanton CA). See **Figure 1**.

**Figure 1.**
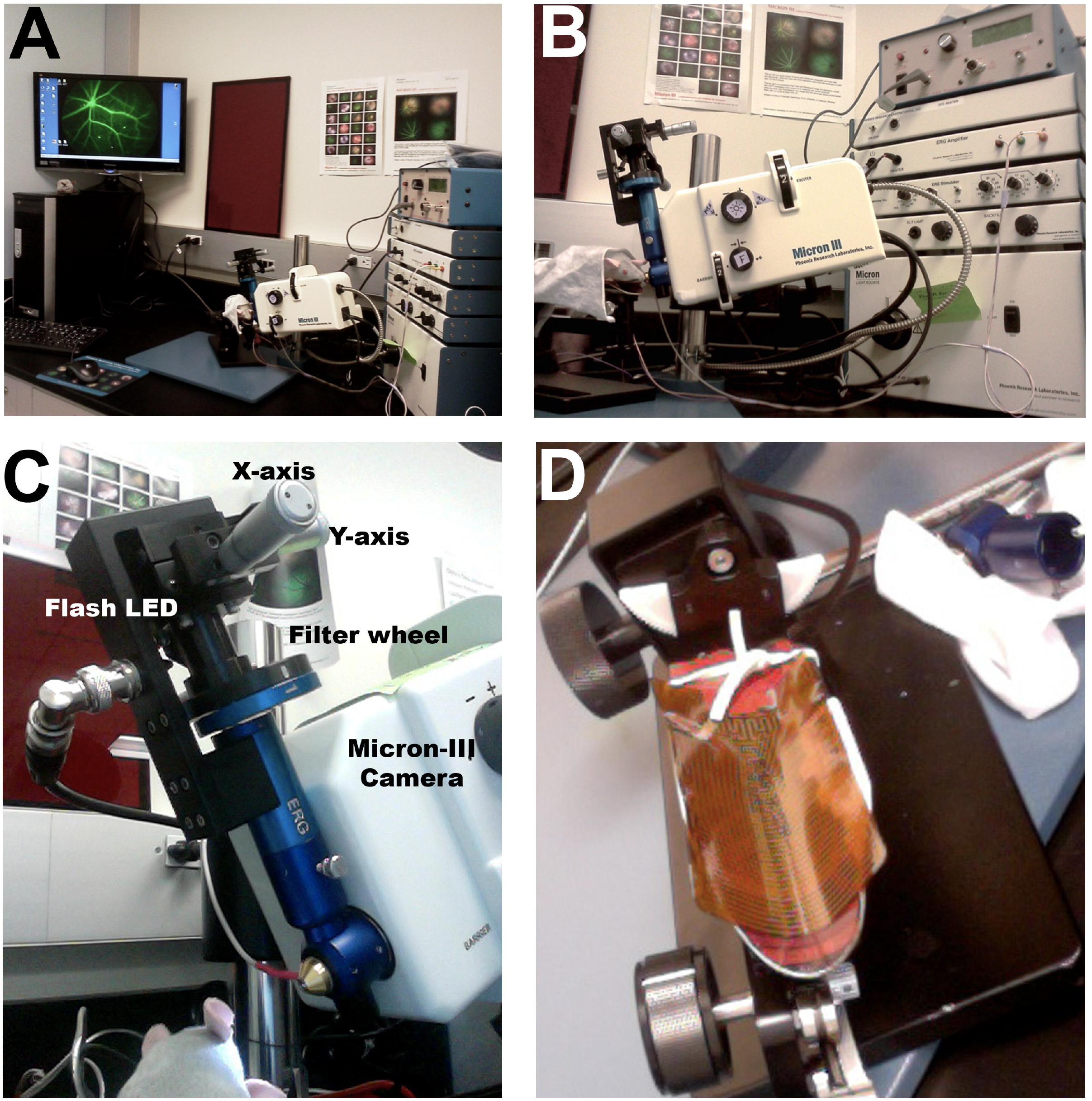
Micron-III System, Pediatric Retinal Research Lab, Eye Research Institute, Oakland University. **A)** Focal ERG work-station. **B)** Micron-III camera with focal-ERG attachment and electrode interface. **C)** Focal-ERG attachment illumination targeting controls. **D)** Mouse warming support.

#### 1. Preparation of retinal imaging and ERG testing suite and required supplies

1. Wipe down the testing room working surfaces with 70% ethanol, as well as the mouse support to be used with the Micron-III (or IV) camera system (Phoenix Research Labs, CA). This is to reduce chances of cross contamination with any murine bacteria or viruses from other colonies or labs that may also use the equipment.
2. Set up a water-based heating pad between 36 to 38° C. This is a recovery station for mice coming out of anesthesia. Small animals have a large surface area/volume ratio and lose their core body heat quickly under anesthesia. Activate the 37° C constant temperature pad on the Micron-III’s mouse support stand. Cover the recovery heating pad and the imaging mouse support pad with saran wrap, which can be discarded later and helps to keep mouse contact surfaces clean.
3. Empty mouse housing bins without bedding should be available, one for each mouse to be tested. Line with paper towel. These are for mice after they awake from anesthesia and prevents them from damaging the cornea surface of their eyes until they are fully recovered and ambulatory.
4. Ensure availability of 1 mL tuberculin syringes, and 26 gage - 1/2 inch single use needles. These will be used for IP injection of anesthesia and for IP injection of 10% fluorescein in PBS.
5. Check that artificial tears, corneal protectant solution (viscous), and ocular petrolatum (petroleum jelly) are available. Artificial tears, over the counter human grade from a pharmacy, are suitable of any brand but select solutions that include calcium, sodium, phosphate, and potassium. Corneal protectant solutions can also be over the counter and are like artificial tears but also have hypromellose or similar polymers to make a thicker gel-like viscosity. These are typically used for human dry-eye treatment, and they provide both hydration but also lubrication. These thicker lubricants will be applied to the mouse cornea for FA-imaging/SD-OCT and focal-ERG as the Micron-III ocular lenses are designed to contact the corneal protectant solution for optimal imaging with maximum field of view of the mouse retina.
6. Check for a sufficient supply of iris-dilation medications, obtained from a veterinary supply company. Two are required, that act on separate muscles systems to cause full dilation of the iris for maximum pupil size: Tropicamide Ophthalmic Solution (5%) USP, and Phenylephrine Hydrochloride Ophthalmic Solution (10%) USP.
7. Check for supply of both Ketamine HCl and Xylazine which are used together for sufficient anesthesia and muscle relaxation. Note that in the United States, researchers must hold a federal DEA license to order, store, and use veterinary grade Ketamine HCl. Depending on the State, researchers may also require a second pharmacy-research license from their State. (This is true in Michigan.) Similar requirements will be in place in other countries.
8. Sterile 10% Sodium Fluorescein solution will be required for Fluorescein Angiography mode imaging.

#### 2. Fluorescein Angiography (FA) Imaging and SD-OCT imaging

Imaging of the retinal vasculature of OIR-treated, or control, mice is combined with SD-OCT image scans of retinal layer thickness to find the locations of retinal zones with different levels of damage or changes. In the case of the OIR-model, the more severely affected retinal zones have experience the most death and loss of inner retinal neurons. Once these areas are mapped in an eye, then a later session with focal-ERG (next section) can be used to compare these different geographic zones for functionality.

1. Set up the Micron camera system with the SD-OCT injector attachment and calibrate the SD-OCT digital scan target indicator line according to the manufacturer’s instructions.
2. Transport mice between assigned housing rooms and the retinal imaging suite while in their normal “shoe box” cages, with food, water, and a filter-cover lid. It is important to transfer mice into fresh cages with fresh bedding.
3. To achieve a brief anesthesia (30 minutes), mice received a single injection (IP) of 50 mg/kg Ketamine HCl and 7 mg/kg Xylazine. The dose is prepared so that 0.005 mL of mixture per gram body weight will provide the final dose required. This also keeps the anesthesia injection small enough to easily deliver in one IP injection.
4. As mice become immobile, drops of artificial tears are added to keep corneas hydrated. Long whiskers around the eyes are trimmed short, or they will interfere with imaging.
5. Give the mouse an injection of 25 uL of 10 % sterile Sodium Fluoresceine (IP).
6. Transfer the mouse to lay on a Kim wipe atop the Micron-III’s constant heat pad (37 °C).
7. The corneal protectant gel solution is applied to both eyes. In addition to preventing corneal dehydration, this thick solution provides optical coupling to the lens of the Micron-III’s SD-OCT attachment.
8. Cover the mouse with a small felt cloth to keep the animal warm during imaging.
9. Starting with the Micron-III SD-OCT camera adjusted about 6 centimeters away from the right eye of the mouse, estimate where the optical axis of the eye is currently aimed. Arrange the mouse using the support stand adjustments for up, down, roll, and swivel to aim the eye towards the SD-OCT lens.
10. Activate the Micron-III camera’s illumination, for bright field mode (white light) and while watching the camera view screen, center the mouse eye in the view and move camera closer to the eye. If the pupil of the eye moves away from center, stop moving the camera closer and adjust mouse position. A few iterations of this process will make it easy for the lens to reach and contact the corneal protection gel on the front surface of the eye. The retina will then fill the camera view. Make small adjustments to center the optic disc in the field of view.
11. Turn the filter selection dial on the Micron-III camera to the fluorescein filter set and select mode to averaging of 15 frames. This will allow for relatively low intensity of illumination and excellent imaging of the retinal microvasculature while selecting areas to capture SD-OCT scans to reveal the retinal layer thicknesses in cross section.
12. Use the SD-OCT target line to position to desired locations on the live FA video image of the retina and capture the desired linear scans (Linear B-scans). Capture video still images with the target guideline in view to keep a record of all B-scan target locations.
13. While imaging one eye, make sure the non-imaged eye is kept protected with fresh application of corneal protectant.
14. When imaging and SD-OCT scans are completed, transfer the mouse to an adsorbent pad and rinse the thicker corneal protected off the eyes with fresh drops of artificial tears.
15. Transfer the mouse to the warming recovery pad and apply ocular petrolatum gel onto the full exposed eye surface. Keep mice dry by changing small pieces of paper towel under their hind leg area as required.
16. As mice wake from anesthesia, and can crawl freely, place them into a recovery tub lined with paper towel (one mouse per tub) until they become fully ambulatory. Then mice can be returned to regular housing.

#### 3. Dark adaptation of mice and preparation for Focal-ERG testing

1. Transport mice between assigned housing rooms and the retinal imaging and ERG testing suite while in their normal “shoe box” cages, with food, water, and a filter-cover lid. It is important to transfer mice into fresh cages with fresh bedding.
2. Cages are set up inside a dark adaptation metal cupboard, that has adequate ventilation. This is within the ERG testing room, equipped dim-red darkroom lighting. After 1.5 hours of dark adaption in the suite’s dark housing station, pupils of mice were dilated with sequential application of tropicamide and then phenylephrine eye drops. Both eye drops are applied again after one minute to make certain full pupil dilation will be obtained.
3. Start the Micron-III focal-erg attachment and camera software and check that those systems are still working.
4. To achieve a brief anesthesia (30 minutes), mice received a single injection (IP) of 50 mg/kg Ketamine HCl and 7 mg/kg Xylazine. The dose is prepared so that 0.005 mL of mixture per gram body weight will provide the final dose required. This also keeps the anesthesia injection small enough to easily deliver in one IP injection.
5. As mice become immobile and will stop blinking, drops of artificial tears are added to keep corneas hydrated. Long whiskers around the eyes are trimmed short, or they will interfere with imaging or ERG.
6. The mouse is transferred to lay on a Kim wipe atop the Micron-III’s constant heat pad (37 °C). The platinum needle reference-electrode is inserted subcutaneous into skin on top of the head, midway between the posterior attachment of the ears. This must not be too far back. A small piece of Scotch brand tape is used to hold the reference-electrode’s wire lead on the back fur, so the needle will not move during manipulations of the mouse. Moving this electrode too far back will pick up electrical activity of the heart’s ventricular contractions in addition to any ERG signal.
7. The platinum needle ground-electrode is inserted into the right-side hind flank skin, and it’s wire lead is also taped onto the fur with Scotch tape.
8. Cover the mouse with a small layer of felt cloth. Make sure the mouse is centered on the support stand.
9. After loss of the blink reflex, tested by artificial tear application, the corneal protectant gel solution is applied to both eyes. In addition to preventing corneal dehydration, this thick solution provides optical coupling to the lens of the Micron’s focal-ERG attachment. This solution also provides electrical coupling to the gold-plated lens-mount, which serves as the measurement-electrode.

#### 3. Aiming and collection of Focal-ERG test data

1. Starting with the Micron-III focal ERG camera adjusted about 6 centimeters away from the right eye of the mouse, estimate where the optical axis of the eye is currently aimed. Arrange the mouse using the support stand adjustments for up, down, roll, and swivel to aim the eye towards the focal-ERG lens. Start about 5-6 centimeters from the eye.
2. The focal-ERG attachment uses its own integral LED light source and has a filter wheel to select white-light or a red-light. Set this filter to red and use the LabScribe control software to activate constant LED illumination, with a relatively dim intensity setting of 9. A second wheel is also dialed to change the aperture for projecting light spots of different size onto the retina. Set the spot size to 1 (largest), for viewing the eye area on the Micron-III camera software.
3. Aiming of the light stimulus is accomplished by viewing the retina with dim red-light illumination using 80% camera gain with 15-frame video averaging. Watching the camera view screen, center the mouse eye in the view and move camera closer to the eye. If the pupil of the eye moves away from center, stop moving the camera closer and adjust mouse position. A few iterations of this process will make it easy for the lens to reach the corneal protectant solution on the front surface of the eye and contact the fluid. The retina will then fill the camera view. On the red field of view, the optic disc will be a small lighter-colored circular zone that is a good landmark for aiming of test flashes. The mouse support can be rolled to tilt the eye up or down as required to center the optic disc in the retinal view.
4. Using previously obtained FA and SD-OCT imaging data, the positions of different retinal geographic locations of interest are first mapped around the disc by noting their specific angle (1-360 degrees, clockwise) and their distance from the edge of the disc measured as measured in disc width equivalents.
5. With the disc centered, adjust the spot aperture (still red-light) to decrease the spot size. In this case to a size as small as the optic disc itself. This will be centered on the optic disc, then use the X, Y axis controls on the focal-ERG head to move the spot to the desired target location.
6. Turn off the constant illumination mode using the LabScribe focal-ERG control software. Set the light-color selection dial to white-light (no red filter) to prepare for white-light flashing.
7. Set flash duration to 30 msec, with full a bright intensity setting. This corresponded to an energy delivery of 43,775 cd-sec/m^2^ of projected retinal area, sufficient to elicit a maximum mixed cone-rod response. The stimulus spot diameter on the retina is about 0.27 mm, area 0.057 mm^2^. The Micron-III focal-ERG had a high luminous efficiency cool-white LZ1-00CW00 LED (LED Engin, San Jose, CA) with emission from 430-650 nm. Triggering of the light stimulus and acquisition of the ERG traces were accomplished with LabScribe2 software equipped with the Phoenix Research Lab ERG module. A setting of twenty stimulus traces were averaged to obtain the average focal-ERG trace, using a 100 msec delay between acquisition cycles. Band pass was set to range from 0.5 Hz to 2000 Hz and the data digitization rate was set to 5000 Hz.
8. When Focal-ERG scans are completed, transfer the mouse to an absorbent pad and rinse the thicker corneal protective gel off the eyes with fresh drops of artificial tears.
9. Transfer the mouse to the warming recover pad and apply ocular petrolatum gel onto the full exposed eye surface. Keep mice dry by changing small pieces of paper towel under their hind leg area as required.
10. As mice wake from anesthesia, and can crawl freely, place them into a recovery tub lined with paper towel (one mouse per tub) until they become fully ambulatory. Then mice can be returned to regular housing.

##### Virtual Microscopy for Morphology Analysis

Enucleated whole eyes were fixed in Davison’s fixative. Fixed tissues were processed for paraffin sections and stained with hematoxylin & eosin. Sections were obtained near the optic nerve region to obtain full cross-sections of retina (7 μm thick) from periphery to periphery. Whole slides were digitized using a 20x objective lens and an Olympus SL120 Virtual Microscopy Slide Scanner and saved in the vsi-file format. Digital files were managed and analyzed using Leica (Slidepath) Digital Image Hub and Digital Slide Box (DSB) web servers, with the Safari web browser (Apple, Cupertino, CA).

## REPRESENTATIVE RESULTS

Focal-ERG data were collected by targeting the same central retinal locations of the same eyes at three different ages: P21, P28 and P42. **Figure 2** illustrates the initial identification of the disc in an eye (left eye). The same spot was then decreased in size, in this case a spot size selection was fixed to a diameter of 0.27 mm, which was just larger than the optic disc itself (**Figure 2B**). This target size was used in continuous red-light mode to visualize placement into the desired target region, relative to the disc. Just prior to acquisition, the focal-ERG system was switched to flash illumination mode and the red filter removed for full white LED stimulation. Then ERG recordings were obtained in regions around the disc in the order: temporal, nasal, superior and inferior to the disc.

**Figure 2.**
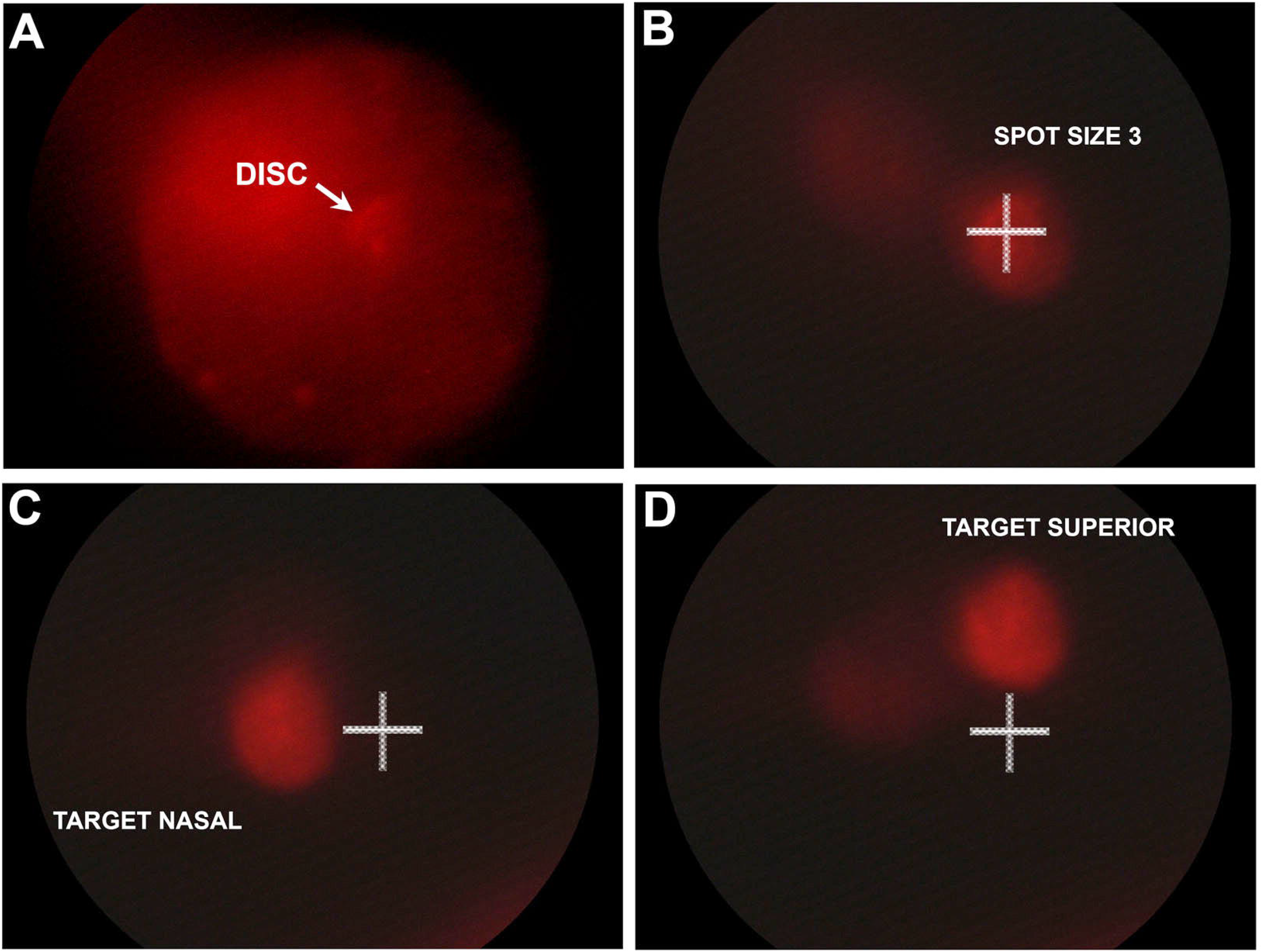
Focal-ERG Targeting Process of the dark-adapted retina. Images shown were obtained during acquisition of the P42 Oxygen-treated focal-ERG data shown in Figure-5. **A)** Dim red-light illumination, not visible to the naked eye, was used to visualize the optic disc by with high camera gain and 15-frame high-speed image summation. **B)** The illumination spot size was then reduced for targeting a circular area of 0.06 mm^2^, just larger than the disc itself. For illustration purposes, the disc location is marked with a white cross. **C)** An example of targeting nasal to the disc, left eye. **D)** Example, targeting superior to the disc, same eye.

By using this targeting scheme, the same central regions (damaged by OIR), could be re-tested, longitudinally, in the same retinas following recovery at ages P21, P28 and P42. A-wave and B-wave amplitudes from the four locations in the retina were averaged at each age compared between normal and control retinas by t-test.

### Loss of inner retinal neurons in the central retina of OIR mice

Examples of retinal morphology are shown in **Figure-3** for control (room air) and OIR mice using the five-day 75% oxygen treatment model at ages P21, P28 and P42. This confirmed that our model was working as expected. By age P21, regions of INL thinning have resulted due to the loss of bipolar cells during the vascular ablation phase in the model. By P21 neovascular growth has restored oxygen to the central inner retinal zones that were ablated. OIR retinas have central regions of INL (inner nuclear layer) thinning of varied severity. These included complete loss of the INL and ganglion cell layer, as well as regions of transition between zones of extensive bipolar cell loss (vascular ablation regions) and zones of less severe cell loss. In these experiments we did not see any significant loss of photoreceptor cells in the OIR mice compared to room air controls. The ONL (outer nuclear layer) of OIR retinas appeared to maintain the same thickness as their normal air counterparts. As expected the OIR treatment mostly impacted the survival of inner retinal neurons that died during the period of hypoxia.

**Figure 3.**
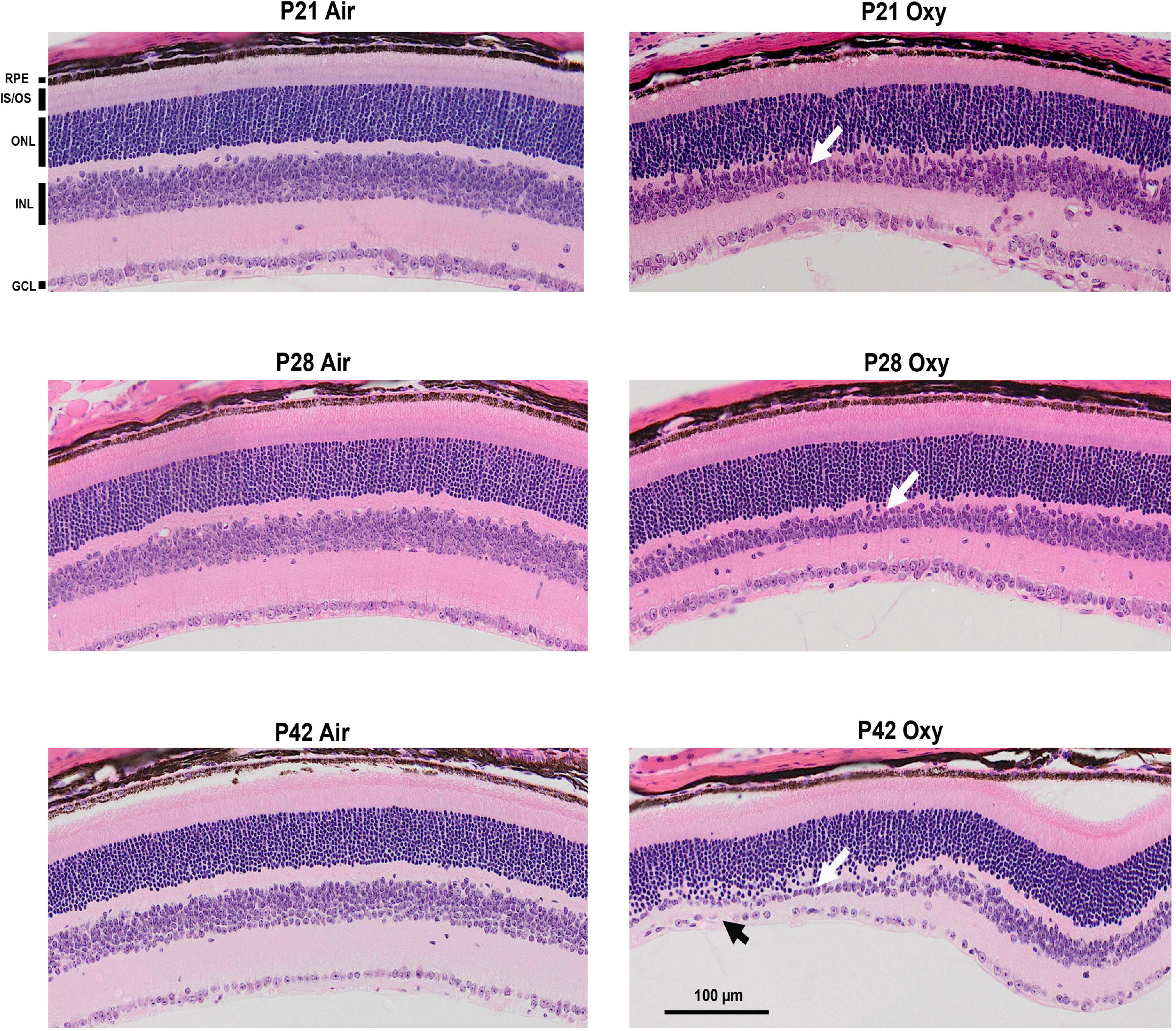
Erosion of the INL and GCL in the central retina of the OIR mouse model. Light microscopy sections of mouse retinas are shown for normal control (room air) and oxygen-treated mice at the same three ages used for focal-ERG recordings, illustrating that ablation of the retinal vasculature during the 75% oxygen treatment results in a varied amount of bipolar cell loss by age P21. Some examples of thinning of the inner nuclear layer and transition to regions of near normal looking retina are shown at ages P21, P28 and P42. Regions of bipolar cell loss are evident (white arrows). Some retinal regions are severely affected to the point where ganglion cell density is also decreased (black arrow). The peripheral retina is generally spared. (RPE - Retinal pigment epithelium; IS/OS – Inner and Outer Segments; ONL – outer Nuclear Layer; INL – Inner Nuclear Layer; GCL – Ganglion cell layer)

### Focal ERG B-wave intensity in areas of different INL thickness

To demonstrate that we can use focal-ERG testing of different small areas within the same retina, we used SD-OCT to map zones of different INL thickness un an OIR retina to select discrete targets for stimulation and recording of the ERG. The loss of bipolar cells and thus INL thinning was more severe in central regions of vascular ablation compared to central regions that do not experience as much vessel ablation. This was confirmed by imaging the same retina on two different days using fluorescein angiography guided SD-OCT. Live image guided SD-OCT shows the retinal layers in a central zone that was ablated of retinal vessels as well as an adjacent region that was not ablated at age P18. (See Figure 4A,C) By age P23, after revascularization, the formerly ablated zone had a relatively thinner INL (Figure 4B,D).

**Figure 4.**
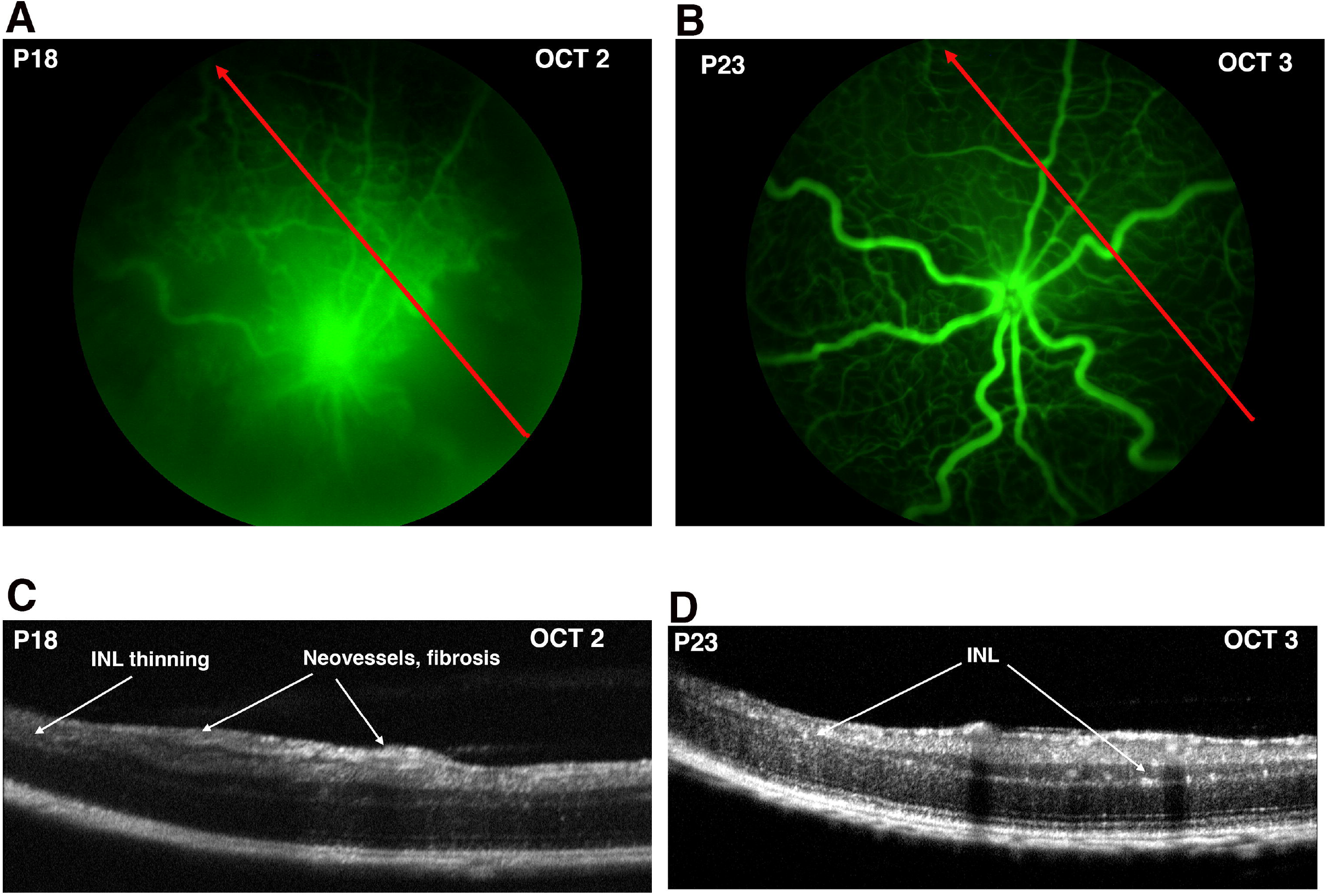
Relation of central avascular zones to INL thinning. INL thinning is more severe in central avascular zones. SD-OCT imaging of the same OIR retina at ages P18 and P23. SD-OCT scans were taken, in the direction of the red arrows, within the period of aggressive neovascularization at age P18 and at age P23 after resolution of vascular regrowth. **A)** At age P18, a linear OCT scan location was selected using fluorescein-angiography guided-imaging to compare an avascular zone (start of scan) transition into a vascular zone. Some fluorescein image image-blur was apparent, from imaging through perfused vessels that are still present on the lens posterior. **B)** The same retina subjected to a repeated OCT linear scan at age P23 after revascularization of the central retina avascular zones. **C)** The OCT image at age P18, corresponding to the location shown above in panel A. **D)** OCT image at age P23, corresponding to the location shown above in panel B.

This same retina was also mapped in more detail at age P23 with several SD-OCT scans to locate central zones with relatively thinner or thicker INL. (Figure 5A) Based on this information the same retina was tested by focal-ERG at age P37 to compare two zones of relatively thinner INL to two zones with relatively thicker INL at the same distance from the disc (Figure 5B).

**Figure 5.**
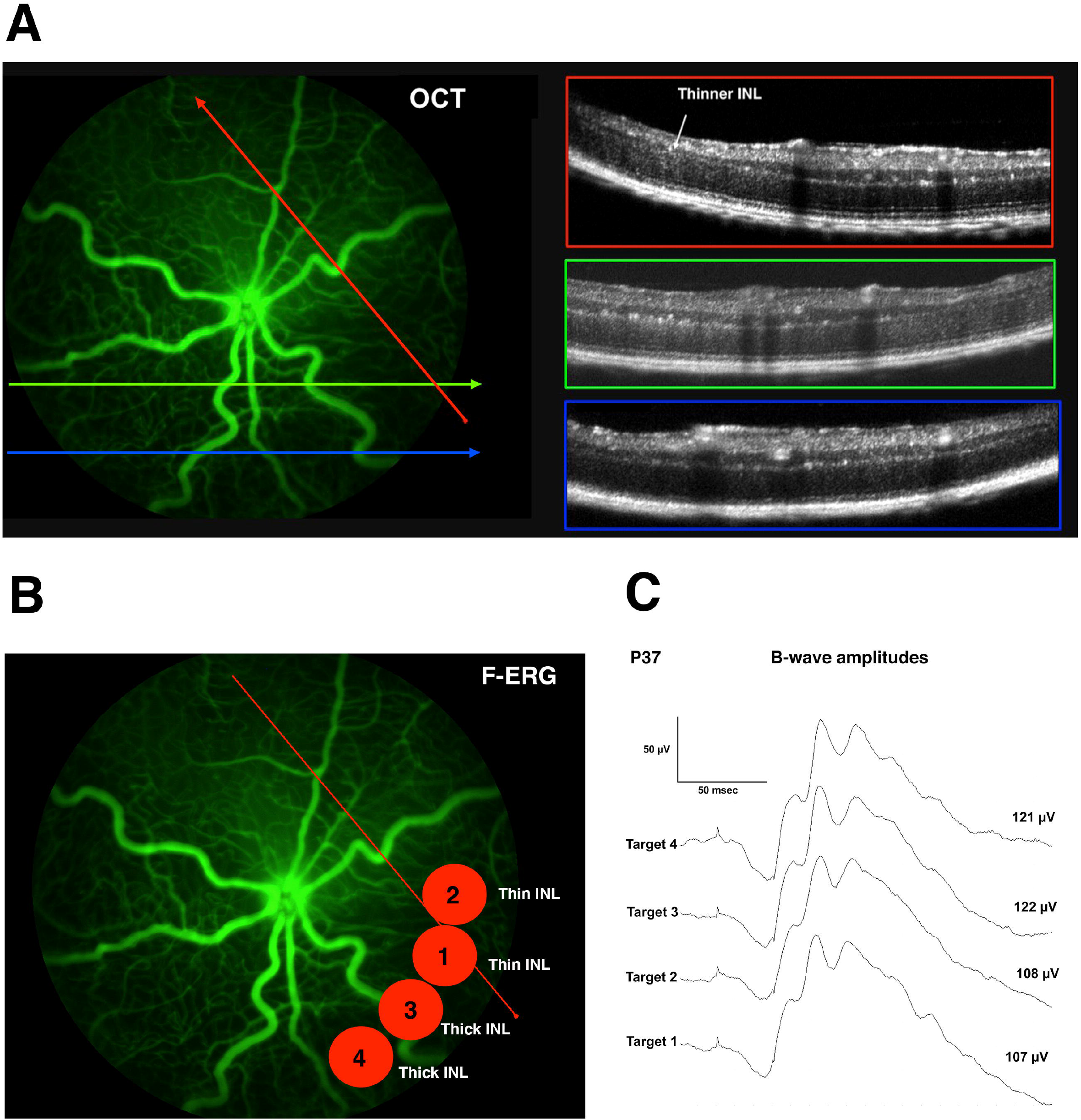
Local variations in B-wave amplitude detected relative to surviving INL thickness. The same retina shown in Figure-3 was mapped by SD-OCT to identify locations of relatively thinner and thicker INL for functional comparison. **A)** To establish that the focal-ERG stimulation can compare B-wave intensity in small adjacent regions with different amounts of bipolar cell loss after oxygen-induced retinopathy (OIR), an OIR retina P23) was first mapped using SD-OCT with linear scans (1.4 mm long). Scans were placed precisely during live imaging with fluorescein angiography. Extensive bipolar cell loss was visible in all OCT scans of the central retina. SD-OCT scan directions are indicated by lines with arrows and the line colors (red, green, blue) correspond to the OCT images in the same-colored boxes to the right. The start of each scan corresponds to the left-side of the OCT image. An example of a relatively thinner INL layer with more extensive bipolar cell loss is indicated (white arrow) in the topmost scan (red). **B)** In a follow up session (age P37) four small circular areas of equal distance from the optic disc were targeted for focal-ERG. Bright flashes were used to elicit a mixed rod-cone response using circular projected light flashes of actual size shown by the numbered red circles (0.27 mm diameter). Targets 1 and 2 were selected to represent regions of relatively thinner INL compared to target regions 3 and 4. **C)** Regions were focal-ERG tested in the relative temporal order 1 to 4. Both regions of relatively thinner and similar INL thickness (targets 1 and 2) had a smaller B-wave amplitude than the relatively thicker regions (targets 3 and 4).

Focal-ERG recordings were made during the bright flash stimulation (mixed rod-cone response) and are shown in Figure 5C. Two target regions with relatively thin inner retinas (labeled 1 and 2) were tested, followed by testing of two regions with relatively thicker inner retinas (labeled 3 and 4). Targets with thinner inner retinas had similar B-wave intensities, which were less than the intensities derived from targets with thicker inner retinas. With the ability to detect differences in B-Wave intensity between regions of different INL thickness that were essentially adjacent, we concluded that any interference at a distance would not preclude focal-erg testing of zones that are much further apart, in completely different retinal quadrants.

### Local recovery of B-wave in central affected regions of the OIR mouse retina

To demonstrate the ability to follow functional changes over time in the same retina we followed an OIR retina and a normal retina over a period of three weeks using image-guided focal-ERG. The same areas of the same retinas were compared longitudinally starting at age P21, after revascularization of the central ablated zone that occurs in this model. The fluorescein angiograms of the retinas followed over three weeks are shown at age P42. (**Figure 6**). This mouse strain had endogenous expression of the YFP protein in a subset of retinal ganglion cells that are also visible prior to injection of fluorescein. Figure 6 shows imaging of the retina through the Focal-ERG’s lens using a fluorescein filter set just after the final ERG testing at age P42. The typical appearance of the normal retinal vasculature was seen in the control retina (Figure 6A). In contrast the OIR retina displayed the torturous retinal vasculature that is familiar and characteristic of retinas that have undergone vascular ablation and neovascular regrowth (Figure 6B).

**Fig 6.**
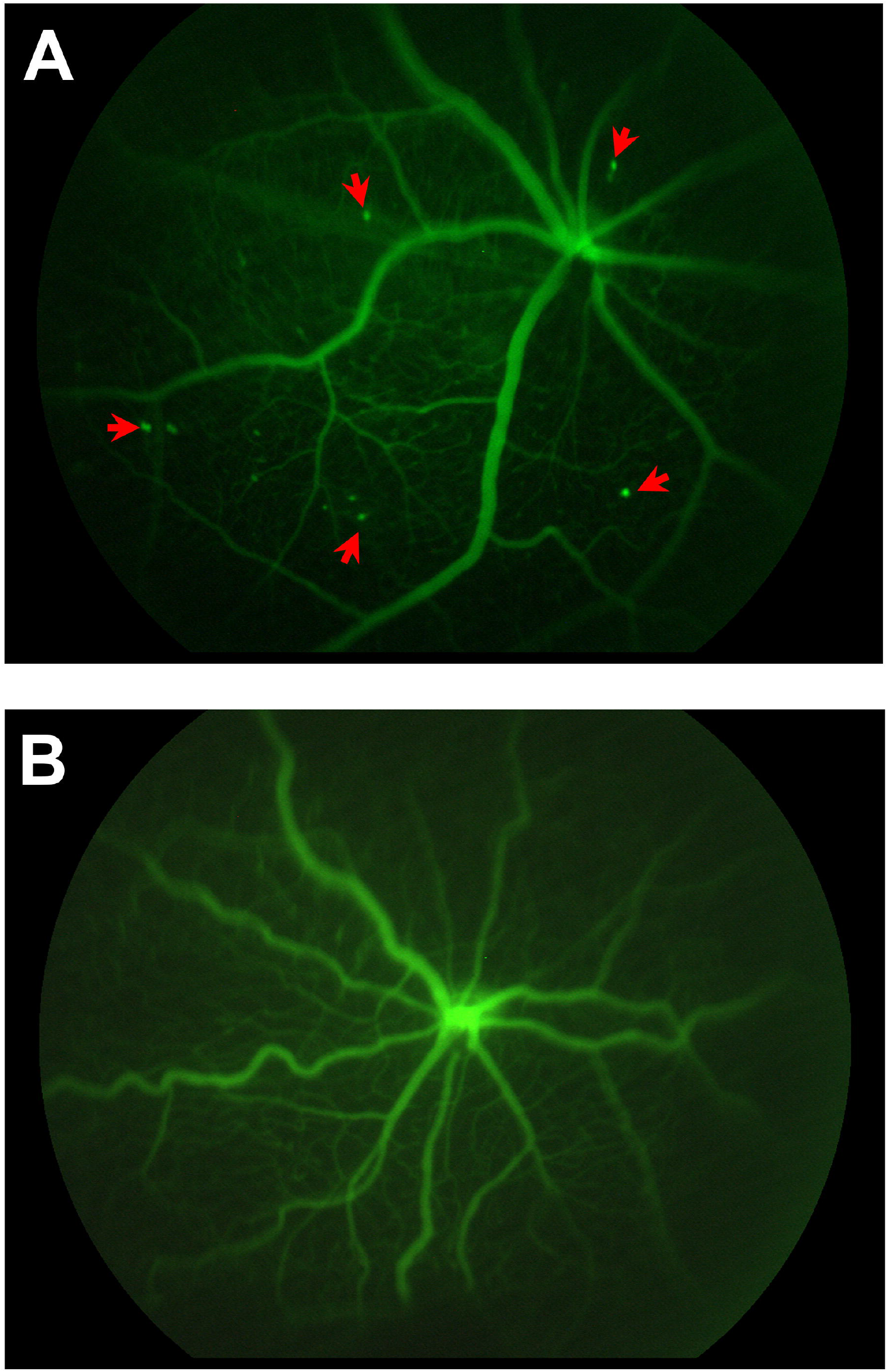
Post-ERG Imaging: Fluorescein Angiography and Ganglion Cells (YFP). **A)** Normal room-air control retina, age P42. In this strain a subset of ganglion cells are also visible (red arrows) from endogenous yellow fluorescent protein using the standard fluorescein filters. Images were captured using the Micron-III’s main light through the focal-ERG lens, immediately after collecting focal-ERG data. **B)** Note the torturous vessel morphology in the OIR retina after neovascularization, age P42.

Four zones were tested, targeting small circular areas adjacent, but not on, the optic disc. Targets were placed superior, inferior, nasal and temporal around the disc. This targeting process facilitated the testing of the same areas at all three ages for each retina: P21, P28 and P42. Focal-ERG traces from the four locations of a normal room air (control) retina and an OIR damaged retina are shown are shown in **Figure-7**.

**Figure 7.**
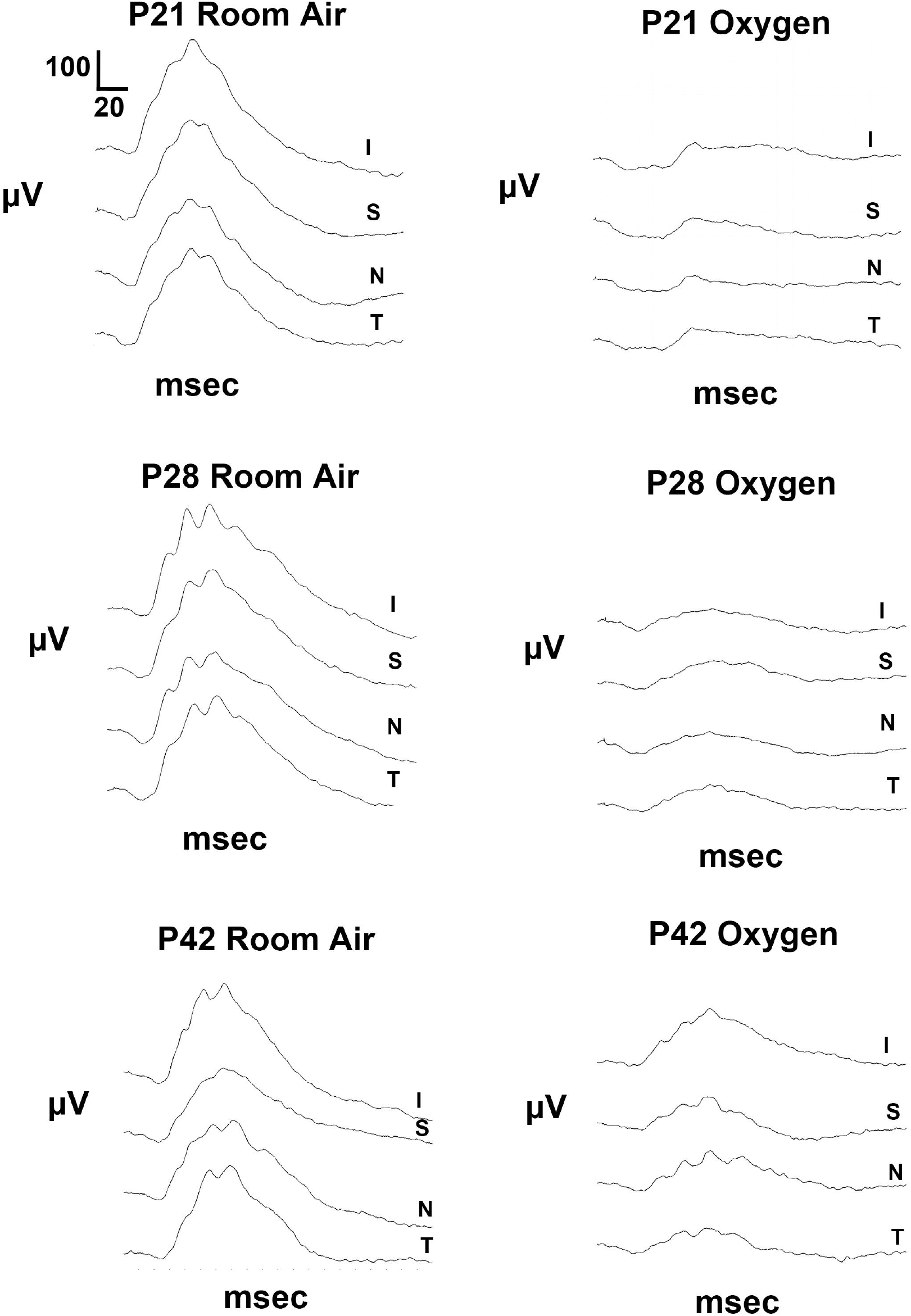
Partial recovery of the B-wave in the OIR damaged neural retina. Focal-ERG traces of the combined rod-cone response were obtained using bright flashes of the same four locations in of the same retinas, followed at three different ages: P21, P28 and P42. The control and OIR retinas were tested in the same sessions on the same day. ERG traces from the four locations around the central retina are shown: superior, inferior, nasal and peripheral (relative to the disc). All four locations in the normal retina displayed a normal looking ERG pattern with A-wave, B-wave and oscillatory potentials. In the OIR damaged retina, at P21 the A-wave could be detected with very little B-wave response. Longitudinal analysis of the same retina areas repeated at ages P28 and P42 revealed a gradual appearance and improvement of the B-wave amplitude in the OIR damaged retina, including some oscillatory potentials.

Just after revascularization at P21, a very week B-wave (average 21.9 mV) was seen in all four tested zones of the OIR damaged retina. In comparison, the focal-ERG B-wave amplitudes of a normal retina at age P21 were 10-fold larger (t-test, P<0.0001). See **Table-1**. The normal retina focal-ERG B-wave traces also displayed familiar oscillatory potentials as normally seen with full-field ERG. (**Figure-7**) The weak focal-ERG B-waves of the OIR retina did not display oscillatory potentials at age P21. In contrast to the different B-wave amplitudes between normal and OIR retinas, the photoreceptor-derived A-wave amplitudes were not significantly different between the control and OIR damaged retinas. (Table 1).

**Table 1:** Focal ERG Average A-wave and B-wave Amplitudes (N=4 locations). ERG response was monitored in the same eyes of normal air (control) and OIR-treated mice at three ages: P21, P28, and P42. Revascularization is completed by P21 in the OIR model and the neural networks in the retina manage to improve the B-wave response about 4-fold from P21 to P42. The B-wave is sub-normal in amplitude because significant areas of the neural retina have permanently lost inner retina neurons (bipolar cells and ganglion cells).

| | A-Wave ($\mu\text{V}$) | | | B-Wave ($\mu\text{V}$) | | |
| --- | --- | --- | --- | --- | --- | --- |
|  | P21 | P28 | P42 | P21 | P28 | P42 |
| Normal Air | $-24.5 \pm 9.2$ | $-26.0 \pm 6.9$ | $-20.1 \pm 4.9$ | $221.7 \pm 37.2$ | $232.2 \pm 27.9$ | $195.6 \pm 29.2$ |
| OIR | $-31.3 \pm 7.7$ | $-22.3 \pm 5.2$ | $-16.6 \pm 4.5$ | $21.9 \pm 16.4$ | $40.4 \pm 13.7$ | $80.9 \pm 37.4$ |
| P-value<br>T-test | 0.15 | 0.21 | 0.16 | <0.0001 | <0.0001 | <0.005 |

Continuing to compare the focal ERG amplitudes at ages P28 and P42, there was a the expected recovery of the B-wave amplitude in the OIR central retinal zones over a three-week span. (**Figure-7**). By age P42 the B-wave amplitudes at each zone tested also began to develop characteristic oscillatory potentials characteristic of retinas with interconnected inner retinal neurons. The local improvement in average focal-ERG B-wave amplitude by age P42 was substantial, about 4-fold greater than that seen at age P21. However, this recovered B-wave amplitude was only 41% of the average B-wave amplitude of an age matched normal retina. In contrast to the lost and recovery of the central retina derived B-wave response, the focal-ERG A-wave amplitudes were not different between the OIR and normal retinas at any of the ages tested.

## DISCUSSION

The mouse OIR model creates ischemic retinal areas in vivo, from ablation of capillary beds, which then triggers a robust revascularization. This model is popular to study the pathophysiology of inner retinal neuron loss during the ischemic period in addition to the regulation of neovascular growth [3,11]. For demonstration purposes here, we have utilized the phase after neovascularization to combine multimodal imaging of the retinal vasculature with function testing of the retina using focal-ERG. With focal-ERG we could limite the light-flash stimulation to extremely small retinal zones that were just larger than the disc. This provided the means to test the function of the inner retina in the OIR-damaged area, based on the ERG B-wave response that is dominated by bipolar cell depolarization [14]. While other human, rat and mouse studies provide good evidence that there is an overall average recovery of post-photoreceptor function [6,8,10], our use of focal-ERG stimulation to longitudinally monitor the same small retinal areas the same OIR retina over a period of several weeks demonstrates how the combination of FA, SD-OCT and Focal-ERG can guide targeting of specific areas to monitor this kind of B-wave recovery.

In the normal mouse retina, the focal-ERG A- and B-wave amplitudes are generally less than when stimulating the entire retinal area (full-field ERG), but the ERG features are familiar and similar in pattern to that of the full-field ERG. A-wave, B-wave and oscillatory potentials were clearly derived and reproducible using the time averaging of twenty traces per area tested. Quite small retinal areas (0.27 mm diameter) were tested close to the disc, to ensure the targeting of central retinal regions that experienced vascular ablation. These same regions experienced bipolar cell loss during the ischemia phase of the model that resulted after 75% oxygen-treated mice were returned to room air (21% oxygen). It is established by other laboratories, and our own, that aggressive neovascular growth returns the inner retina’s blood supply by age P21 [3,12,13]. With the substantial death of bipolar cells during the ischemic phase, we demonstrate here that the recovery of an initially lost B-wave response can be monitored as it recovers after day P21 in this model. Revascularization results in recovery from ischemia and the potential for remaining retinal neurons to regain some functionality.

Our histology done after confirmed the loss of neurons seen in SD-OCT is not an artifact we concluded that most of the cell death in the OIR damaged central retinal zones involved the inner retina, seen as a reduction in the INL thickness. In contrast the ONL remained a normal thickness (10-12 photoreceptor nuclei) even in regions where almost all bipolar neurons were lost (Figure 3). This reflects the fact that photoreceptor cell inner-segments, and their oxygen-demanding mitochondria, are adjacent to the RPE/choroid and the choroid’s blood supply. Thus, photoreceptors cells are less reliant on the three vascular beds of the inner retina, which were ablated during the 75% oxygen treatment phase. This expectation was consistent with the near normal A-wave amplitude seen in OIR damaged central retinal zones when compared to room-air control central retinal zones. Photoreceptors remained normal in number and functionally mature enough to respond to light stimulation and generate a negative A-wave of similar amplitude to the normal retina.

Consistent with the loss of bipolar cells in the OIR-model, the central retinal B-wave amplitudes were substantially reduced and abnormal in their pattern at age P21 in the OIR retina. At P21 the B-wave was essentially decimated. Furthermore, longitudinal follow-up of the same local central zones at ages P28 and P42 revealed a progressive and significant recovery of the B-wave amplitude. The recovery of familiar oscillatory potentials superimposed on the B-wave was also apparent in the OIR focal-ERG by age P42.

Multiple modes of analysis using imaging of the retinal vasculature (FA), retinal layer thickness (SD-OCT) and focal-ERG can provide an improved understanding of the localized effects of retinal vascular damage on retinal neuron survival and functionality in vivo. This kind of analysis can be used to monitor the potential loss, and also recovery, of function over time (i.e. synaptogenesis) and also a metabolic recovery from pan-retinal changes in energetics and ion regulation [15].

## ACKNOWLEDGMENTS

Research funding from the Pediatric Retinal Research Foundation, (Virginia and Clarence Clohset estate) to KPM and KD. Oakland University Center for Biomedical Research to KPM. Special Trustees of Moorsfields Eye Hospital, National Institute of Health Biomedical Research Center at Moorsfields Eye Hospital, UCL Institute of Ophthamalology, TFC Frost Trust, HCA International, and Moorsfields Surgeons Association for visiting fellowship to SCW. Authors acknowledge Oakland University student Jennifer Felisky, for assistance in manuscript file preparation and reviewing.

## DISCLOSURES

None.

## Notes

### Competing Interest Statement

The authors have declared no competing interest.

### Summary of Updates

This version of the manuscript has been revised to show the methodology of combined focal ERG with Fluoresceine Angiography and SD-OCT for structural-functional analysis of the retinal neurovasculature. It is written in a format for method guidance.

## REFERENCES

1. Gole GA, Browning J, Elts SM. The mouse model of oxygen-induced retinopathy: a suitable animal model for angiogenesis research. Doc Ophthalmol 1990; 74:163–9.

2. Ricci B. Oxygen-induced retinopathy in the rat model. Doc Ophthalmol 1990; 74:171–7.

3. Smith LE, Wesolowski E, McLellan A, Kostyk SK, D’Amato R, Sullivan R, D’Amore PA. Oxygen-induced retinopathy in the mouse. Invest Ophthalmol Vis Sci 1994; 35:101–11.

4. Krohne TU, Westenskow PD, Kurihara T, Friedlander DF, Lehmann M, Dorsey AL, Li W, Zhu S, Schultz A, Wang J, Siuzdak G, Ding S, Friedlander M. Generation of retinal pigment epithelial cells from small molecules and OCT4 reprogrammed human induced pluripotent stem cells. Stem Cells Transl Med 2012; 1:96–109.

5. Fulton AB, Hansen RM, Moskowitz A, Akula JD. The neurovascular retina in retinopathy of prematurity. Prog Retin Eye Res 2009; 28:452–82.

6. Fulton AB, Hansen RM, Moskowitz A. The cone electroretinogram in retinopathy of prematurity. Invest Ophthalmol Vis Sci 2008; 49:814–9.

7. Akula JD, Hansen RM, Martinez-Perez ME, Fulton AB. Rod photoreceptor function predicts blood vessel abnormality in retinopathy of prematurity. Invest Ophthalmol Vis Sci 2007; 48:4351–9.

8. Fulton AB, Akula JD, Mocko JA, Hansen RM, Benador IY, Beck SC, Fahl E, Seeliger MW, Moskowitz A, Harris ME. Retinal degenerative and hypoxic ischemic disease. Doc Ophthalmol 2009; 118:55–61.

9. Harris ME, Moskowitz A, Fulton AB, Hansen RM. Long-term effects of retinopathy of prematurity (ROP) on rod and rod-driven function. Doc Ophthalmol 2011; 122:19–27.

10. Nakamura S, Imai S, Ogishima H, Tsuruma K, Shimazawa M, Hara H. Morphological and functional changes in the retina after chronic oxygen-induced retinopathy. PLoS One 2012; 7:e32167.

11. Pierce EA, Avery RL, Foley ED, Aiello LP, Smith LE. Vascular endothelial growth factor/vascular permeability factor expression in a mouse model of retinal neovascularization. Proc Natl Acad Sci U S A 1995; 92:905–909.

12. Wang L, Shi P, Xu Z, Li J, Xie Y, Mitton K, Drenser K, Yan Q. Up-regulation of VEGF by retinoic acid during hyperoxia prevents retinal neovascularization and retinopathy. Invest Ophthalmol Vis Sci 2014; 55:4276–87.

13. Tokunaga CC, Mitton KP, Dailey W, Massoll C, Roumayah K, Guzman E, Tarabishy N, Cheng M, Drenser KA. Effects of anti-VEGF treatment on the recovery of the developing retina following oxygen-induced retinopathy. Invest Ophthalmol Vis Sci 2014; 55:1884–92.

14. Saszik SM, Robson JG, Frishman LJ. The scotopic threshold response of the dark-adapted electroretinogram of the mouse. J Physiol 2002; 543:899–916.

15. Berkowitz BA, Roberts R. Evidence for a critical role of panretinal pathophysiology in experimental ROP. Doc Ophthalmol 2010; 120:13–24.

